# Trichoplein controls endothelial cell function by regulating autophagy

**DOI:** 10.1101/595702

**Authors:** Andrea Martello, Angela Lauriola, David Mellis, Elisa Parish, John C Dawson, Lisa Imrie, Martina Vidmar, Noor Gammoh, Tijana Mitic, Mairi Brittan, Nicholas Mills, Neil O Carragher, Domenico D’Arca, Andrea Caporali

**Affiliations:** University/BHF Centre for Cardiovascular Science, QMRI, University of Edinburgh, Edinburgh UK; Department of Biomedical, Metabolic and Neural Sciences, University of Modena & Reggio Emilia, Modena, Italy; Cancer Research UK Edinburgh Centre, Institute of Genetics and Molecular Medicine, University of Edinburgh, Edinburgh UK; Centre for Synthetic and Systems Biology (SynthSys), University of Edinburgh, Edinburgh UK

**Keywords:** Autophagy, endothelial cells, pericentriolar material, autophagosome maturation, SQSTM1/p62, GABARAP.

## Abstract

Autophagy is an essential cellular quality control process that emerged critical for vascular homeostasis. Here we describe, the role for Trichoplein (TCHP) protein in linking autophagy with endothelial cells (ECs) function. The depletion of TCHP in ECs impairs migration and sprouting. TCHP directly binds PCM1, to regulate degradation of GABARAP, thus leading to a defective autophagy. Mechanistically, TCHP is indispensable for autophagosome maturation and its depletion resulted in the accumulation of SQSTM1/p62 (p62) and unfolded protein aggregates in ECs. The latter process is coupled to TCHP-mediated NF-kB activation. Of note, low levels of TCHP and high p62 levels were detected in primary ECs from patients with coronary artery disease. In addition, *Tchp* knock-out mice showed accumulation of p62 in the heart and cardiac vessels and reduced cardiac vascularization. Here, we reveal an autophagy-mediated mechanism for TCHP down-regulation, which poses a plausible target for regulation of endothelial function.

## Introduction

Autophagy is an essential quality control function for the cell to maintain its homeostasis, through selectively degrading harmful protein aggregates and/or damaged organelles. This is a vital intracellular process for recycling nutrients and generating energy for maintenance of cell viability in most tissues and in adverse conditions (Galluzzi et al, 2017a). Basal autophagy mediates proper cardiovascular function (Bravo-San Pedro et al, 2017). Varity of cardiovascular risk factors can cause defective autophagy in vascular cells, producing high levels of metabolic stress and impairing functionality of endothelial cells (ECs) (De Meyer et al, 2015). Autophagy has been shown to regulate angiogenic activity and the release of von Willebrand factor from ECs (Torisu et al, 2013). In addition, endothelial specific deficiency of autophagy is pro-inflammatory and pro-senescent, as it promoted atherogenic phenotype in a murine model of atherosclerosis (Vion et al, 2017).

Specific autophagic receptors are responsible for selective autophagy by tethering cargo to the site of autophagosomal engulfment (Stolz et al, 2014). The recognition of ubiquitinated substrates is provided by molecular adaptors including p62/SQSTM1 (p62), which bind on one side to ubiquitin and, on the other end, to autophagosome-specific proteins (like members of the LC3/GABARAP/Gate16 family). The interaction between p62 and LC3/GABARAP bridges the autophagic machinery with its cargo, thereby fostering the selective engulfment by the autophagosome (Yu et al, 2018). In mammalian cells, six ATG8 orthologues exist that are divided into the LC3 and GABARAP subfamilies. It was reported that LC3 and GABARAP function non-redundantly during autophagosome biogenesis. Specifically, LC3 subfamily members promote elongation of phagophore membranes (Weidberg et al, 2010), whereas GABARAP is critical in the closure of the phagophore membrane (Tsuboyama et al, 2016) and fusion of autophagosomes with lysosomes (Nguyen et al, 2016). Recent studies demonstrated that a pool of GABARAP exist in the centrosome and peri-centrosomal region and regulate autophagosomes formation during amino acid starvation (Joachim et al, 2017).

The levels of p62 are regulated transcriptionally and through continuous degradation during basal autophagy. The defective autophagy, however, induces accumulation of p62, followed by the formation of aggregates (Komatsu et al, 2007). This is further observed in human ECs in the Cerebral Cavernous Malformations disease (Marchi et al, 2015), and in human smooth muscle cells whereby p62 accumulation accelerated the development of stress-induced premature senescence (Grootaert et al, 2015). Beside its role in autophagy, p62 is a scaffolding hub for the cellular signalling pathways involving NF-kB activation, nerve growth factor signalling, and caspase activation (Sanchez-Martin & Komatsu, 2018).

Trichoplein (TCHP) is a cytosolic ubiquitously expressed 62 kDa protein identified as a keratin filament binding protein (Nishizawa et al, 2005). So far, the function of TCHP is deemed dependent on its partner proteins and their cellular localization. In proliferating cells, TCHP serves as a scaffold protein not only for appendage-associated Ninein, involved in microtubule anchoring at the mother centriole (Ibi et al, 2011), but also for the centriole-associated Aurora kinase A activity, implicated in the destabilization of cilia (Inaba et al, 2016). Alternatively, in differentiated, non-dividing epithelial cells, TCHP translocates from the centrioles to keratin filaments, and desmosomes (Nishizawa et al, 2005). TCHP was reported to reside on outer mitochondrial membrane (OMM), where it binds Mitofusin2 (Mfn2), regulating the ER– mitochondria tethering and promoting mitochondria fission (Cerqua et al, 2010; Vecchione et al, 2009). Moreover, increased levels of TCHP enable decorin evoked mitophagy (Neill et al, 2014).

It is still unknown what role TCHP plays in EC function, particularly regarding its localization and mechanisms of action. We report here an unexpected correlation between the loss of TCHP and defective autophagy in ECs. This subsequently leads to accumulation of p62 and an aberrant protein aggregates build-up within ECs, thus inhibiting ECs function.

## Results

### Lack of TCHP impairs endothelial cell function

To examine the impact of TCHP on endothelial function, we performed a Matrigel tubule formation assay. TCHP down-regulation (**Figure S1A**) severely affected the tubule forming capacity of HUVECs *in vitro* (Figure 1A) and the formation of vessels *in vivo* using Matrigel plugs (Figure 1B). In agreement, reduced level of TCHP also affects ECs migration as measured in the wound healing (Figure 1C). To further dissect the phenotype of ECs lacking TCHP, we analysed the expression of a subsets of genes controlling angiogenesis, inflammation and cell cycle. The expression of IL1β, IL6, IL8, MCP1, CDKN2A/p16 and CDNKNB/p14 was differentially up-regulated upon TCHP down-regulation at 7 days post lentiviral transduction (Figure 1D). Moreover, TCHP knock-down cells exhibited a senescent associated phenotype as seen by the increase of CDKN2A/p16 (**Figure S1B**), β-Galactosidase activity (β-Gal) (**Figure S1C**), and the accumulation of aggregosomes (**Figure S1D**) at 7 days post lentiviral transduction.

**Figure 1:**
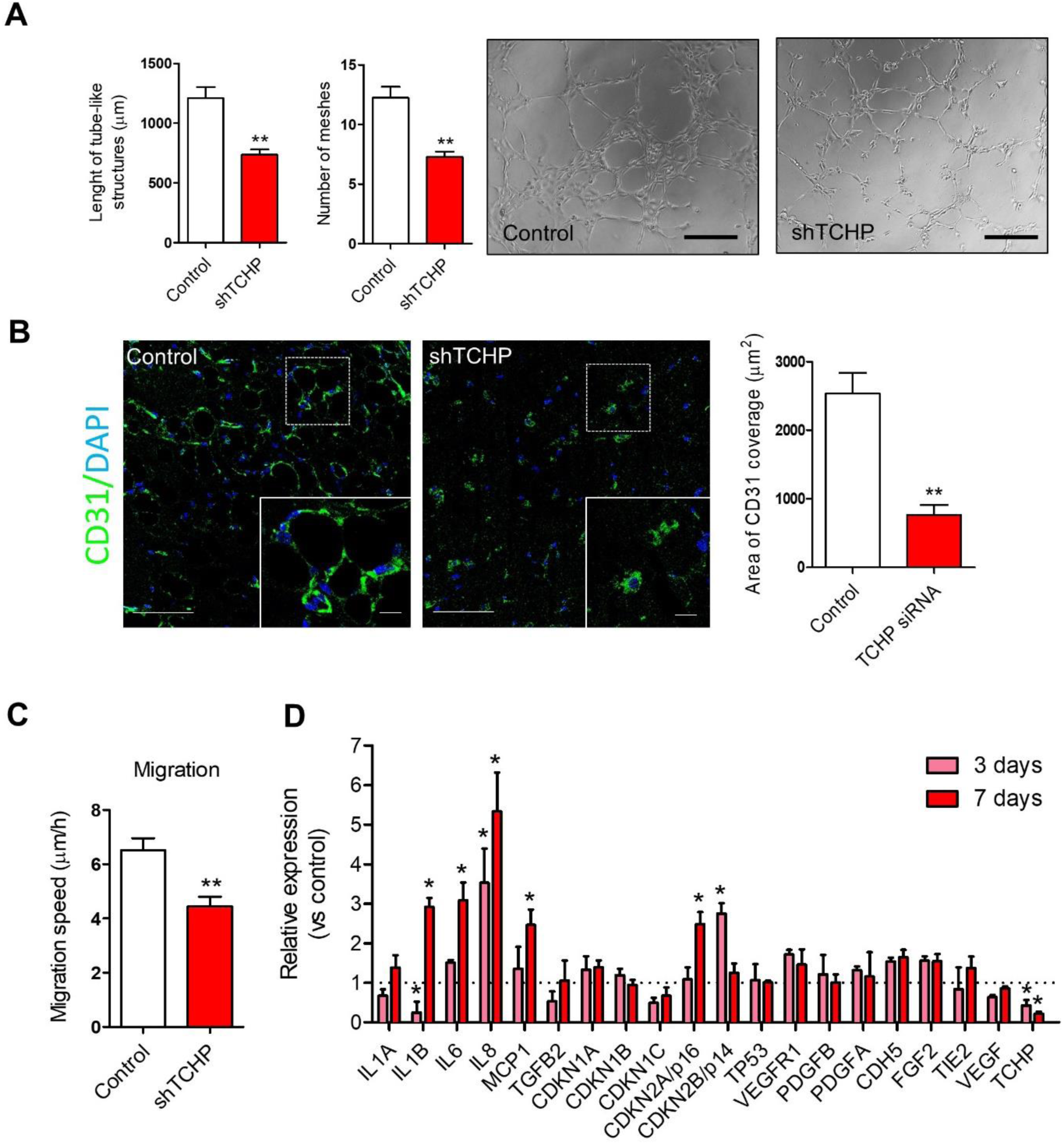
TCHP knock-down affects endothelial cells function. **A**, Endothelial network formation on Matrigel was analysed by quantification of the total length of tubule-like structures and number of meshes; *right panels*, representative microphotographs from Matrigel assay; Scale bar: 100 μm. (n= 5, unpaired t test, **p ≤ 0.01 vs control). **B**, *Right panel*: quantification of the area of CD31 coverage in the Matrigel plugs mixed with TCHP siRNA or control oligos at 21 days after implantation; (n= 5 animals per group, unpaired t test, **p ≤ 0.01 vs control). *Left panels*: representative images showing the new microvessels positive for CD31 (green), in the implanted plugs. Scale bars, 50 µm and 25µm for the inset. **C**, Effect of TCHP knock-down on HUVEC migration speed measured by electric cell-substrate impedance sensing (ECIS) (n=5, unpaired t test, **p ≤ 0.01 vs control). **D**, Relative mRNA levels of TCHP and subset of genes. Graphs represent transcripts measured at 3 and 7 days post TCHP shRNA transduction (n=3, unpaired t test; *p ≤ 0.05 vs control). For panel A, B, C, and D: Data are represented as mean ± SEM

### TCHP binds PCM1 directly to regulate its localization

To identify TCHP interacting partners, we performed co-immunoprecipitation (Co-IP) coupled with mass spectrometry analysis using FLAG-tagged TCHP as a bait in HEK293 cells. The centriolar satellite protein PCM1 was identified as the most enriched protein in anti-FLAG pull-down in comparison with the control experiment (Table 1). We validated the mass spectrometry results showing that in HEK293 cells two different FLAG-tagged version (N- and C-terminal) of TCHP co-immunoprecipitated with endogenous PCM1 (Figure 2A). TCHP displays two coiled-coil regions, at the N terminus (1–136 AA), which are necessary and sufficient for centriolar localization and function (Inoko et al, 2012). Using FLAG-tagged TCHP deletion mutants, we found that the residues corresponding to the second coiled-coil region (41-136 AA) are critical for the binding to PCM1 (Figure 2B). Moreover, we demonstrated that the interaction between TCHP and PCM1 is conserved in HUVECs, since TCHP is co-immunoprecipitated with endogenous PCM1 (Figure 2C).

**Table 1:**
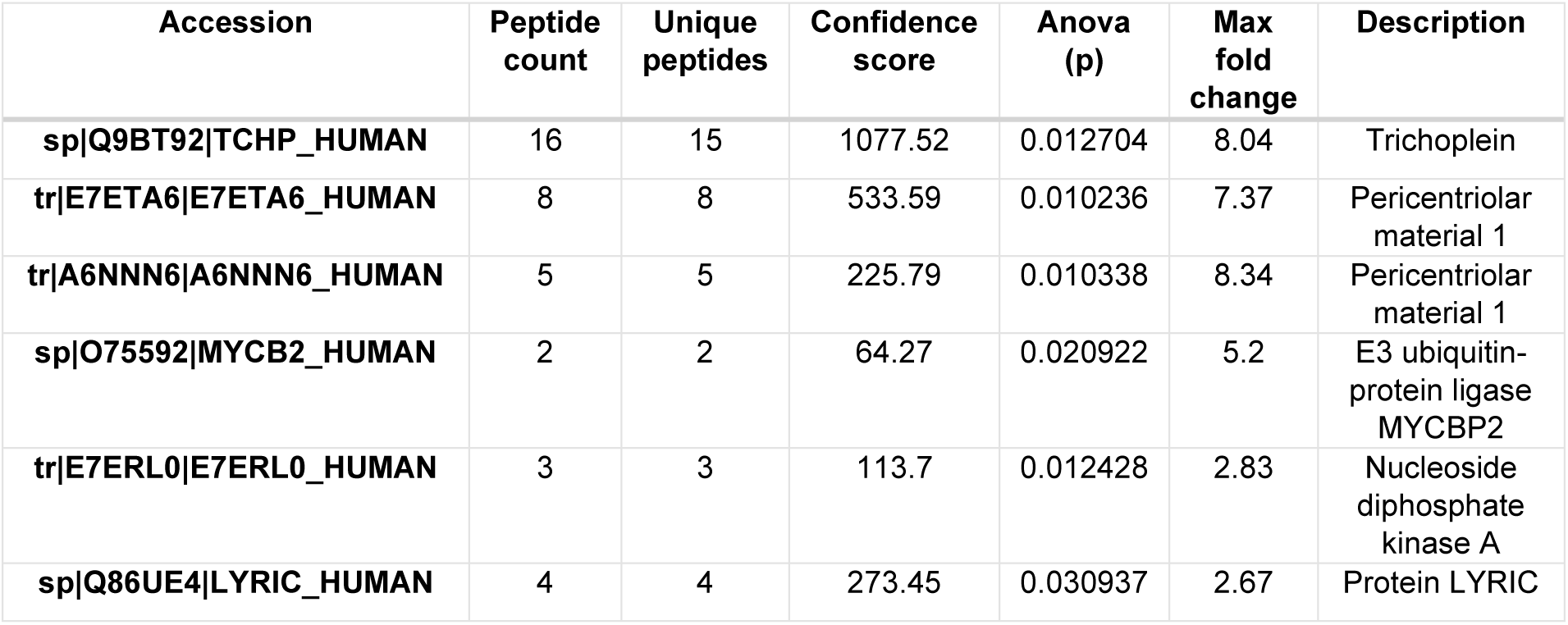
Mass spectrometry analysis of proteins co-immunoprecipitated with TCHP-FLAG in HEK293 cell extracts.

**Figure 2:**
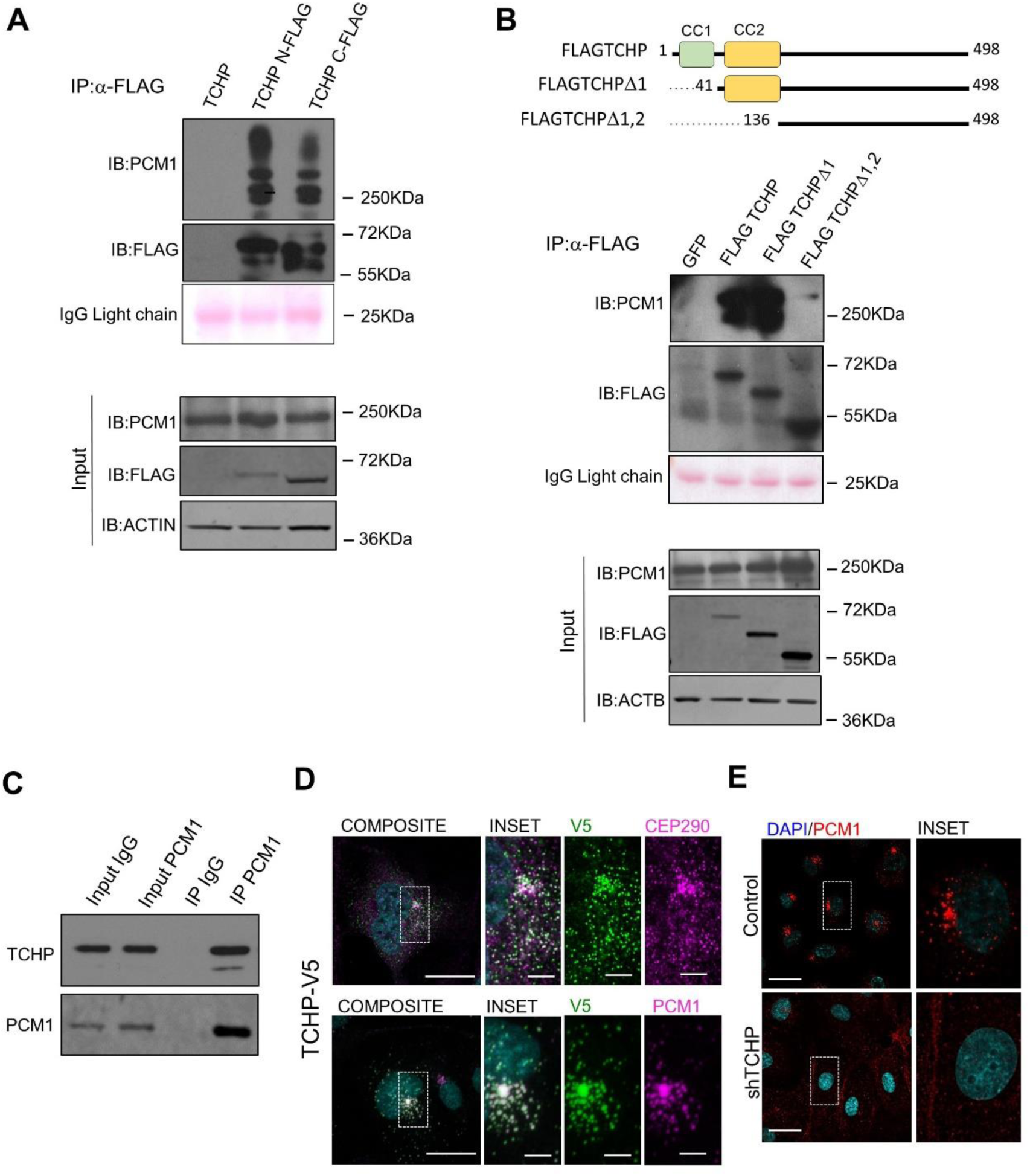
TCHP interacts directly with PCM1. **A**, Immunoblot analysis showed HEK293 transfected for 48 hours with expression vectors for TCHP, and FLAG-TCHP at N- and C-terminal. **B**, Immunoblot analysis showed HEK293 transfected for 48 hours with expression vectors for TCHP-FLAG C-terminal or constructs with deletions of the coiled coil domain 1 (TCHP Δ1) and 2 (TCHP Δ2), as indicated in the scheme. For **A** and **B**: total lysates were immunoprecipitated with anti-FLAG antibodies, and blots were probed sequentially with anti-PCM1, anti-FLAG and anti-ACTIN antibodies. IgG light chains are indicated with Red Ponceau staining. The input totals were analyzed by parallel immunoblotting as control for the level of expression. **C**, Anti-PCM1 immunoprecipitation from HUVECs cells and TCHP and PCM1 immunoblot. **D**, Co-localization of TCHP-V5, PCM1 or CEP290. HUVECs were transduced with TCHP-V5 lentivirus and stained for anti-PCM1, CEP290 and anti-V5 antibodies. Scale bars, 25 µm and 5 µm in the inset. **E**, TCHP knock-down or control cells were stained for anti-PCM1 antibody. Scale bars, 50 µm and 5 µm in the inset.

We next tested the localization of PCM1 and TCHP in ECs. TCHP displayed a predominately punctate cytoplasmic pattern and concentrates close to the nucleus in the same compartment occupied by the centriolar satellite protein CEP290 (Kim et al, 2008). We established that TCHP extensively co-localized with PCM1 in the pericentriolar matrix and satellite region (Figure 2D). Interestingly, depletion of TCHP had significant effect on PCM1 localization, showing a lack of PCM1 accumulation at the centrosome (Figure 2E).

### TCHP regulates PCM1 and GABARAP stability

Since constant turnover of PCM1 is regulated by the proteolytic degradation (Didier et al, 2008), we analyzed PCM1 protein levels and degradation rate in the TCHP-depleted cells. Like centrosomes and cytoskeleton-associated proteins (Sedjai et al, 2010), PCM1 was enriched in the Triton X-100 insoluble fraction and TCHP depletion reduced PCM1 at the steady state (Figure 3A). Next, we used cycloheximide (CHX) and MG132 to block the translation and proteasomal degradation, respectively. In TCHP depleted cells treated with CHX, PCM1 degradation was enhanced compared to control cells, while inhibited by MG132, thus revealing the augmented proteasome-dependent turnover of PCM1 (Figure 3A).

**Figure 3:**
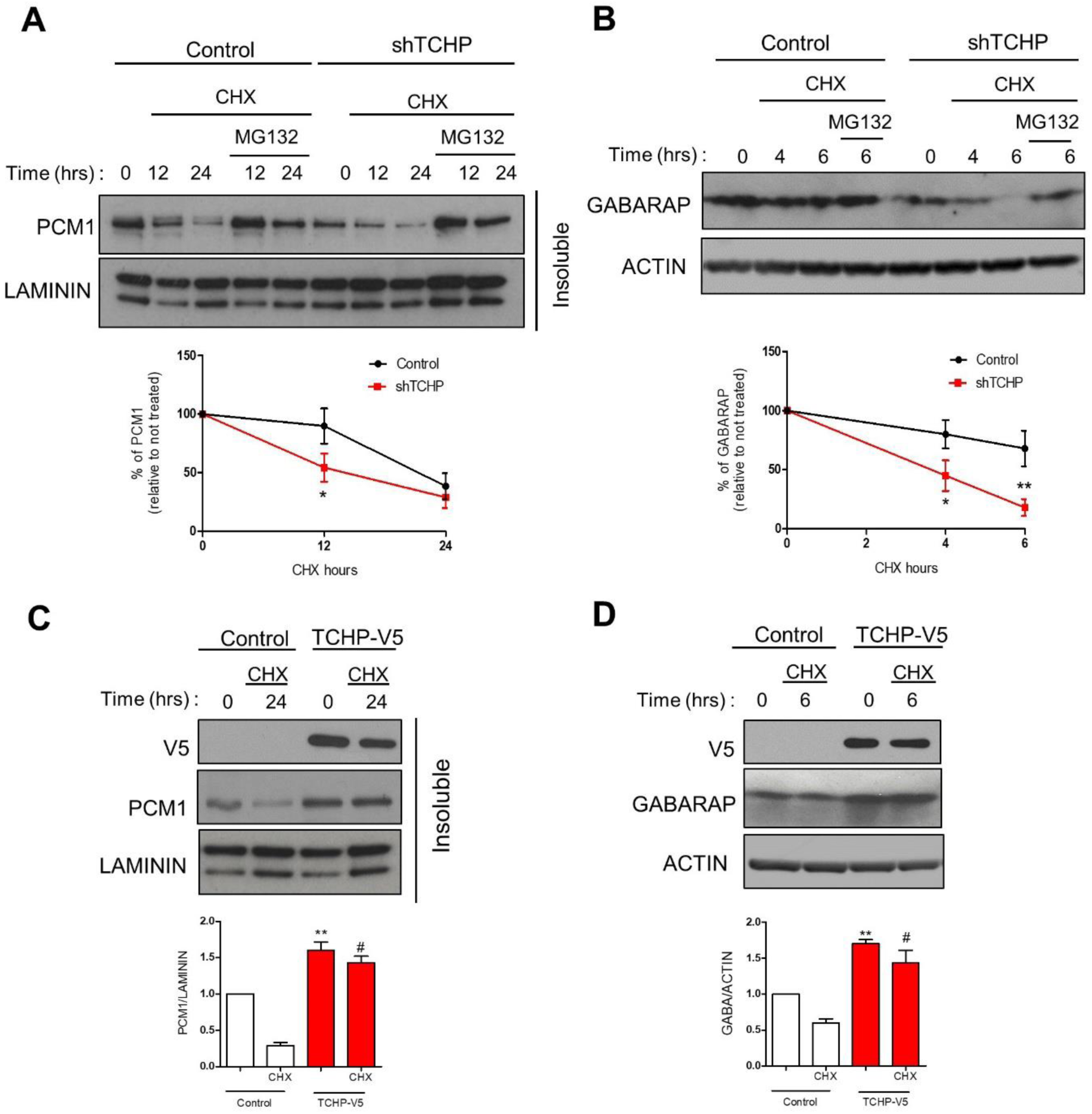
TCHP regulates PCM1 stability. **A**, Control or TCHP knock-down HUVECs were subjected to cycloheximide (CHX) and MG132 treatment for the indicated number of hours prior to immunoblotting. PCM1 levels were analysed in the insoluble fraction. Laminin has been used as loading control. *Below panel*: Quantification of PCM1 degradation. Data are represented as mean ± SEM (n = 3; one-way ANOVA; ∗p ≤ 0.05 vs control time 0). **B**, in the above condition, Western blot for anti-GABARAP and anti-ACTIN antibodies. *Below panel*: Quantification of GABARAP degradation. (n = 3; one-way ANOVA; ∗p ≤ 0.05 vs control time 0). **C**, Control or TCHP overexpressing HUVECs were subjected to CHX treatment for the indicated number of hours prior to immunoblotting. Laminin has been used as loading control. Below panel: Quantification of C. Data are represented as mean ± SEM (n = 3; one-way ANOVA; **p ≤ 0.01 vs control time 0; ^#^p ≤ 0.05 vs control CHX). **D**, in the above condition, Western blot was probed for anti-GABARAP and anti-ACTIN antibodies. Below panel: Quantification of D. (n = 3; one-way ANOVA; **p ≤ 0.01 vs control time 0; ^#^p ≤ 0.05 vs control CHX).

Recent data suggests that GABARAP binds directly to PCM1 and the loss of PCM1 results in destabilization of GABARAP via proteasomal degradation, thus effecting autophagy (Joachim et al, 2017). In the control cells, we confirmed a proteasome-dependent reduction of GABARAP during the 6 hours treatment with CHX. Without TCHP, in line with a reduced stability of PCM1, the rate of GABARAP degradation was enhanced. (Figure 3B). Conversely, ectopic expression of TCHP-V5 increased expression of PCM1 (Figure 3C) and GABARAP (Figure 3D) and extended their stability. Altogether, these data suggest that TCHP regulates GABARAP by proteasomal degradation.

### TCHP down-regulation impairs autophagic homeostasis

Importantly, we next set to establish what role TCHP plays in autophagy. Transmission Electron Microscope (TEM) revealed a significant increase in the number of autophagic vesicles when TCHP is depleted in HUVECs (**Figure S2A**). Alongside reduction in GABARAP, immunocytochemistry staining revealed an increased number of LC3 and p62 positive puncta (Figure 4A). We next set to determine the regulation of the autophagic flux by analyzing the levels of p62 and LC3 in basal and Hank’s buffered (HBSS) starved cells with or without Bafilomycin (BafA1). In full medium, Western blot analysis confirmed an increased level of the lipidated form of LC3 (LC3II) band, an increase of p62 protein levels in TCHP knock-down cells compared with control (Figure 4A).

**Figure 4:**
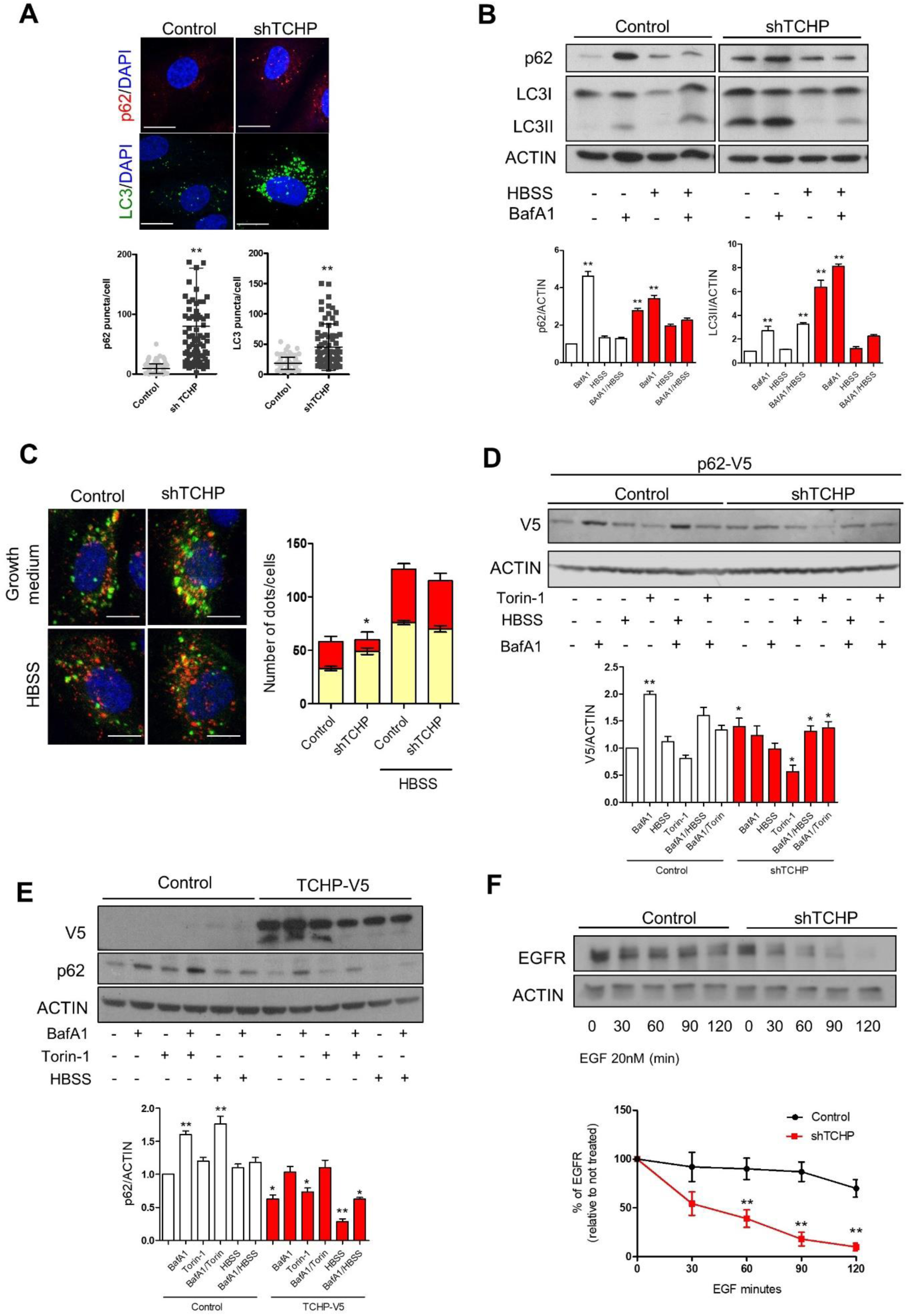
Analysis of autophagy in TCHP-depleted endothelial cells. **A**, Immunofluorescent staining for LC3 and p62 in TCHP knock-down and control cells. Scale bar, 25μm. (n=80, unpaired t test; *p ≤ 0.05 vs control). **B**, Western blot of p62 and LC3 during under normal culture condition or starved condition (HBSS) or the presence of BafA1 (2 hours) in TCHP knock-down or control cells. *Lower panel:* quantification of C, (n = 3; one-way ANOVA; *p ≤ 0.05 vs control time 0). **C**, HUVECs were transduced with the tandem mCherry-EGFP-LC3 and with shRNA TCHP or control vectors. *Left panels*: Representative picture of mCherrry-EGFP-LC3 reporter. Scale bar, 25μm. *Right panels*: Quantification of the number of mCherry-only (red bars, autolysosomes) or double-positive (mCherry^+^/EGFP^+^; yellow bars, autophagosomes). (n=80, three independent experiments; one-way ANOVA; *p ≤ 0.05 vs control). **D**, TCHP knock-down or control cells were transduced with p62-V5 vector. Western blot for anti-V5 antibody during under normal culture condition or HBSS or in the presence of BafA1 or Torin-1 for 2 hours. *Lower panel:* quantification of D, (n = 3; one-way ANOVA; **p ≤ 0.01; **p ≤ 0.05 vs control time 0). **E**, HUVECs were transduced with TCHP-V5 vector. Western blot for anti-V5 and anti-p62 antibody during under normal culture condition or HBSS or in the presence of BafA1 or Torin-1 for 2 hours. Lower panel: quantification of E, (n = 3; one-way ANOVA; **p ≤ 0.01;*p ≤ 0.05 vs control time 0). **F**, Representative Western blot showing Epidermal Growth Factor Receptor (EGFR) levels during pulse chase experiment with EGF (20nM) for the indicated times. *Lower panel:* quantification of E, (n = 3; one-way ANOVA; **p ≤ 0.01 vs control time 0). For panel A, B, C, D, E and F: Data are represented as mean ± SEM.

When autophagic flux was blocked with BafA1 at basal conditions, there was a greater accumulation of p62 in the control cells compared with that in TCHP knock-down cells, while this difference was blunted after autophagy induction. Lipidated LC3II accumulated in TCHP knock-down cells at basal condition. In contrast, the lipidation rate increased more in control cells than in the TCHP knock-down cells after the blockage of autophagy. Finally, the treatment with HBSS re-activated the autophagic flux in TCHP knock-down cells as demonstrated by strong degradation of LC3II and reduction of p62 (Figure 4B).

This data was corroborated when using mCherry-EGFP-LC3 assay as a complementary approach (Kimura et al, 2007). The mCherry fluorescence was lower in TCHP knock-down cells compared with the control, attesting to a decrease in autolysosome formation and a slower autophagic flux in cells lacking TCHP (Figure 4C). There was not a significant difference in the percentage or total number of mature autolysosomes following starvation in HBSS medium (Figure 4C). We also analysed the autophagic flux using V5-tagged p62 (p62V5) as an exogenous substrate of autophagy. At the steady-state conditions in full growth medium, more p62V5 has accumulated in the knock-down compared with control cells. Following BafA1 treatment under full growth conditions p62V5 accumulated in the control cells more than in the knock-down cells (Figure 4D), hence confirming a delay in the autophagic flux in the absence of TCHP. The rate of degradation of p62V5 by activation of autophagy during nutrient starvation or treatment with torin-1 was comparable between TCHP knock-down and control cells (Figure 4D). Finally, overexpression of TCHP reduced the basal level of p62 in ECs and the accumulation of p62 after BaFA1 treatment under full growth medium (Figure 4E). In addition, exogenous TCHP increase degradation of p62 during nutrient starvation or treatment with torin-1(Figure 4E).

The abnormalities affecting the autophagic flux in TCHP knock-down cells suggest a block during terminal stage of autophagy, when autophagosomes fuse with the late endosome or lysosome (Shen & Mizushima, 2014). Immunostaining demonstrated a discrete alteration in shape, distribution and intensity of RAB11, EEA1 and RAB7 vesicles (**Figure S2B**). In addition, knock-down of TCHP altered the cellular positioning of lysosomes as shown by LAMP2 immunolabeling, inducing a marked perinuclear clustering of these organelles. Further, increased intensity of LysoTracker Red™ in TCHP knock-down cells, suggested an increased acidification and activity of lysosomes (**Figure S2C**), hence we performed epidermal growth factor (EGF) receptor (EGFR) degradation assay (Pinilla-Macua & Sorkin, 2015). Endocytosis and subsequent lysosomal-mediated degradation are the major regulators of EGFR stability following ligand activation. Cell stimulation with EGF upon knock-down of TCHP resulted in an increasing degradation of EGFR compared with control (Figure 4F), thus ruling out lysosomal dysfunction as the cause of decreased autophagic flux.

### The depletion of TCHP inhibits autophagosome maturation and efficient delivery of p62 to the lysosomes

Having observed that TCHP knock-down reduces autophagic flux without affecting the degradative capacity of lysosome, we hypothesized that autophagosome maturation could be impeded, thus affecting autolysosome formation. Since GABARAP is critical for autophagosome expansion and maturation (Tsuboyama et al, 2016) (Nguyen et al, 2016), we performed the proteinase K protection assay (Hasegawa et al, 2016) to assessed the efficiency of cargo receptor loading during autophagosome biogenesis and maturation. Autophagic vesicles were isolated by cytoplasm differential centrifugation and treated with proteinase K to determinate the proportion of the cargo receptors p62 and NDP52 not accessible to the protease because protected within autophagosome. We found that the sensitivity of p62 and NDP52 to proteinase K was enhanced by TCHP knock-down in low-speed pellet (LSP) (Figure 5A); however, p62 associated cargo are not sequestered in sealed autophagosome in a greater fraction than NDP52 cargo. Collectively, these results demonstrated the role for TCHP in autophagosome maturation. Furthermore, we analysed the structure of autophagosomes in control and TCHP knock-down cells treated with BafA1 (Figure 5B). Accordingly with documented roles for ATG8s and in particular GABARAP family in enlarging autophagosomal membranes (Nakatogawa et al, 2007), we observed a considerable reduction for the average size of autophagic vesicles (AVs) and autophagosomes (APs) in TCHP Knock-down cells (Figure 5B).

**Figure 5:**
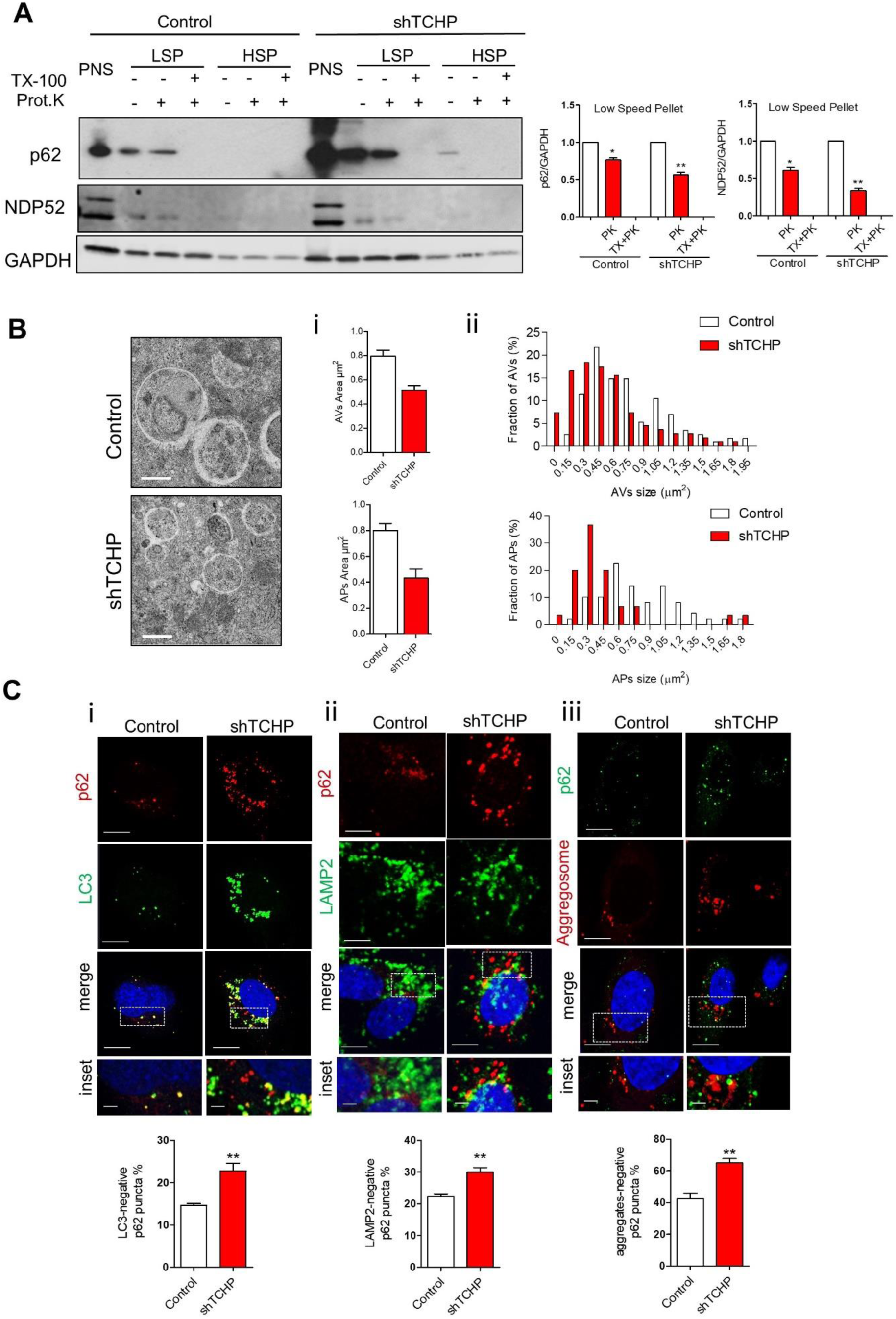
TCHP regulates autophagosome maturation. **A**, The postnuclear fraction (PNS) from TCHP knock-down and control HUVECs was separated into (low-speed pellet) LSP and (high-speed pellet) HSP fractions and then analysed by immunoblots using anti-p62 and anti-NDP52 antibodies. The sub-fractions were treated with proteinase K (Prot. K) with or without Triton X-100 (TX-100). The NDP52 and p62 levels were quantified and normalized to the respective non-treatment control. GAPDH has been used as loading control (n=3 per group; one-way ANOVA, **p ≤ 0.01). **B**, Representative TEM images of autophagosomes in control and TCHP knock-down HUVECs in complete medium after 3h incubation with BafA1 (scale bar 600nm): **i**, quantification of mean autophagic vessicles (AVs) and autophagosome (APs) area and **ii**, distribution of the cross-section areas of the analysed vesicles expressed in percentage. Data in ii and iii Data are represented as mean ± SEM. (n=7 cells; >100 vesicles per sample; unpaired t test **p <0.01 vs control. **C**, Co-localization between **i**, LC3 and p62 **ii**, LAMP2 and p62 and **iii** aggregates and p62 in TCHP knock-down and control cells. Scale bar 25 µm and 2 µm in the inset. *Lower panels*: quantification of B, C and D (n=80, unpaired t test; **p ≤ 0.01 vs control). For panel A, B, C and D: Data are represented as mean ± SEM.

We then examined the subcellular distribution of p62 in ECs stained for LC3 and LAMP2. Knock-down of TCHP increased the percentage of p62 puncta that were also negative for LC3 and LAMP2 (Figure 5C). This suggests that TCHP could impair the efficient clearance of p62 by autophagosomes (LC3-negative p62 puncta) and consequently its delivery to lysosome (LAMP2-negative p62 puncta). The co-localization of p62 with misfolded proteins has already been described in autophagy (Komatsu et al, 2007), however, in TCHP knock-down cells, p62 failed to co-localize with the aggregates (Figure 5C).

### NF-kB inhibition reduces p62 accumulation, restoring endothelial cell function

To elucidate the mechanisms behind the accumulation of p62 in ECs, we have developed a phenotypic screening assay to select compounds, which could decrease the accumulation of p62 in ECs lacking TCHP (Figure 6A and 6B). We screened a library of 176 target-annotated compounds (including protease, epigenetics and kinase inhibitors) at two different doses (0.3 and 3.3 μM) with the primary outcome to reduce the number of p62 cytoplasmic puncta. The screen generated a list of 25 hits (Table 2) with the top ones being excluded due to high toxicity (Apicidin and Terreic acid) or negative effect on endothelial function (Splitomicin) (Breitenstein et al, 2011). The screen identified BAY11-7082 (IKK inhibitor), TYRPHOSTIN AG1288 (Tyrosine kinases inhibitor) and SB202190 (p38MAPK inhibitor) as having a clear effect on reducing p62 accumulation in TCHP knock-down cells (Figure 6C); named compounds were further tested in secondary functional assays. BAY11-7082 was the only compound that was able to reduce cytokine transcription (Figure 6D) and restore migratory capacity in TCHP knock-down ECs (Figure 6E). Since BAY11-7082 was able to effect cells lacking TCHP, we next investigated the activation of NF-kB in these cells. TCHP knock-down cells exhibited a marked increase in NF-kB S536 phosphorylation compared to the control cells. Treatment with BAY11-7082 reduced NF-kB phosphorylation and p62 expression, revealing a NF-kB-dependent contribution to p62 accumulation (Figure 6F). Moreover, TCHP knock-down ECs showed an increased enrichment of NF-kB/p65 on IĸBα (as positive control locus for NF-kB translocation (Sun et al, 1993)) and p62 promoters by chromatin immunoprecipitation (Figure 6G).

**Table 2:** Analysis of compounds reversing p62 accumulation in TCHP knock-down cells

| **Chemical Name** | **Target** | **[Drug]**<br>**uM** | **Cell**<br>**Count** | **Cytoplasmic**<br>**Granule**<br>**Count** | **Prediction**<br>**Score** |
| --- | --- | --- | --- | --- | --- |
| **Splitomicin** | SIRT-2 inhibitor | 3.3 | 211 | 0.53 | 0.84 |
| **Apicidin** | HDAC inhibitor | 3.3 | 170 | 0.42 | 0.79 |
| **BAY 11-7082** | IKK pathway | 0.3 | 237 | 0.36 | 0.78 |
| **SB-202190** | p38 MAPK | 0.3 | 267 | 0.39 | 0.72 |
| **Terreic acid** | BTK | 3.3 | 174 | 0.50 | 0.70 |
| **TYRPHOSTIN AG 1288** | Tyrosine kinases | 3.3 | 215 | 0.31 | 0.69 |
| **Aminoresveratrol sulfate** | SIRT1 activator | 0.3 | 206 | 0.42 | 0.68 |
| **Daidzein** | Negative control for Genistein | 0.3 | 199 | 0.46 | 0.68 |
| **Epigallocatechin gallate** | MMPs | 3.3 | 220 | 0.50 | 0.66 |
| **N-Ethylmaleimide** | Cysteine proteases | 3.3 | 233 | 0.59 | 0.61 |
| **LY 294002** | PI 3-K | 3.3 | 118 | 0.46 | 0.57 |
| **E-64-C** | Calpain; Cathepsins; Papain | 0.3 | 246 | 0.53 | 0.57 |
| **Calpain Inhibitor II (ALLM)** | Calpain; Cathepsins L, B | 0.3 | 254 | 0.63 | 0.56 |
| **Tranylcypromine hemisulfate** | Lysine demethylase inhibitor | 3.3 | 208 | 0.60 | 0.53 |
| **Daidzein** | Negative control for Genistein | 3.3 | 196 | 0.65 | 0.53 |
| **GW 5074** | cRAF | 3.3 | 212 | 0.65 | 0.53 |
| **TYRPHOSTIN 46** | EGFRK, PDGFRK | 3.3 | 203 | 0.54 | 0.52 |
| **Lavendustin A** | EGFRK | 3.3 | 215 | 0.71 | 0.52 |
| **Nullscript** | Scriptaid Neg control | 3.3 | 177 | 0.72 | 0.50 |
| **Suramin-6Na** | SIRT1 inhibitor | 3.3 | 218 | 0.64 | 0.47 |
| **Roscovitine** | CDK | 0.3 | 219 | 0.65 | 0.46 |
| **Z-PLG-NHOH** | MMPs | 0.3 | 250 | 0.60 | 0.45 |
| **Piceatannol** | Syk | 0.3 | 273 | 0.76 | 0.44 |
| **ML-9Å-HCl** | MLCK | 0.3 | 241 | 0.68 | 0.43 |
| **SB-203580** | p38 MAPK | 3.3 | 269 | 0.81 | 0.42 |

**Figure 6:**
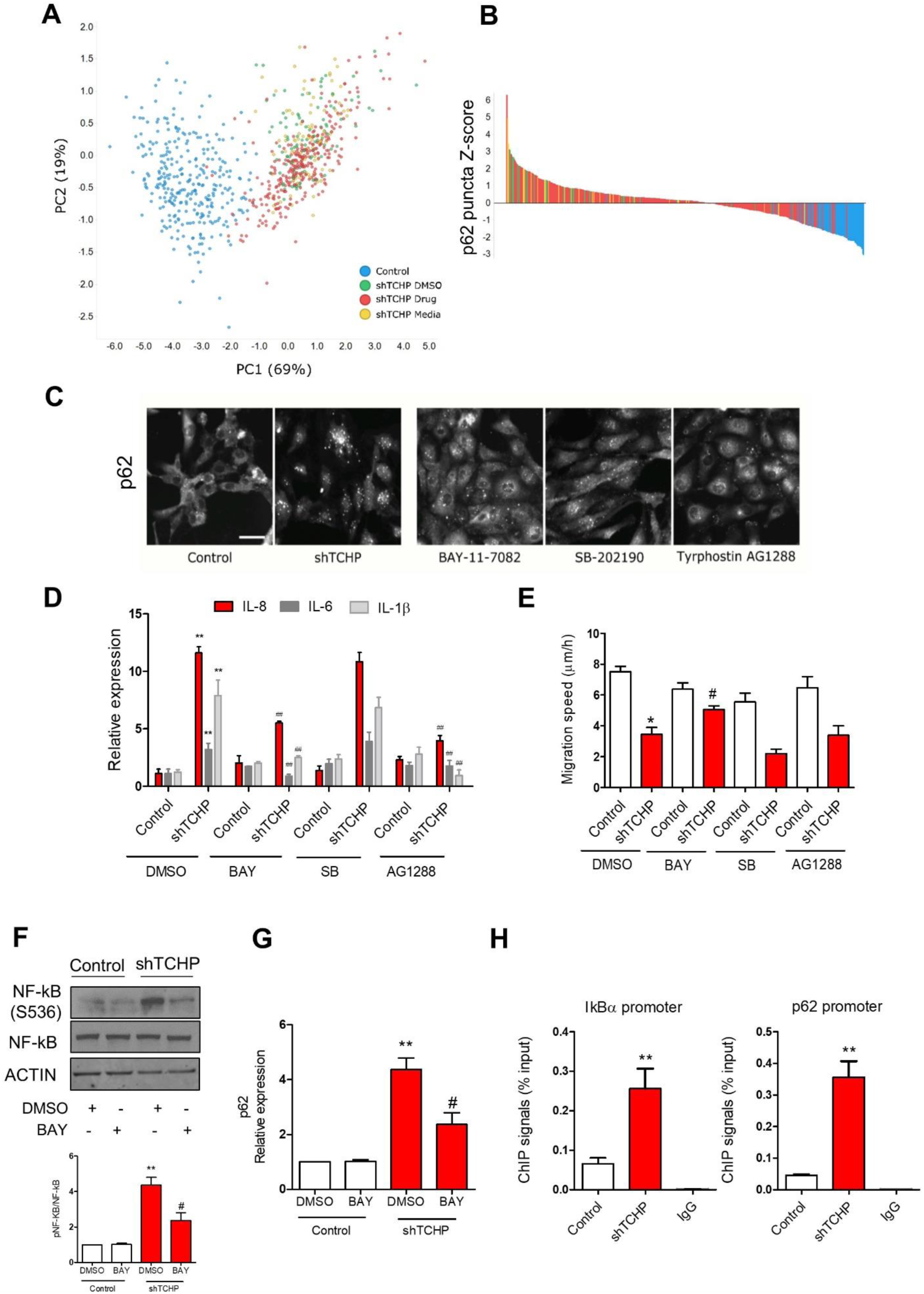
Identification of mechanism of p62 accumulation. **A**, Principal component analysis of compound screen. Principal components (PC) 1 and 2 are show and their respective variance indicated. **B**, Z-score ranking of cytoplasmic puncta of p62 immunostaining. **C**, Representative images for p62 of control and TCHP knock-down HUVECs treated with 3 hit compounds. Scale bar, 50 µm. TCHP knock-down and control cells were treated with BAY11-7082 or TYRPHOSTIN AG1288 or SB202190 or vehicle (DMSO) for 48 hours. **D**, Expression of IL-6, IL-8 and IL-1β and **E**, migration speed was measured; (n=3; unpaired t test; *p ≤ 0.05; **p ≤ 0.01 vs control DMSO; ^#^p ≤ 0.05; ^##^p ≤ 0.01 vs shTCHP DMSO). **F**, Western blot analysis for anti-phospho NF-kB (S536), total NF-kB, in TCHP knock-down and control cells treated with BAY or vehicle. *Below panel:* quantification of F, (n=3; unpaired t test; *p ≤ 0.05; **p ≤ 0.01 vs control DMSO; ^#^p ≤ 0.05; ^##^p ≤ 0.01 vs shTCHP DMSO). **G**, p62 expression in TCHP knock-down and control cells treated with BAY or vehicle. (n=3; unpaired t test; *p ≤ 0.05; **p ≤ 0.01 vs control DMSO; ^#^p ≤ 0.05; ^##^p ≤ 0.01 vs shTCHP DMSO). **H**, ChIP-qPCR analysis confirms the of NF-kB p65 enrichment to IkBα and p62 promoter in TCHP knock-down cells. (n=3; unpaired t test; **p ≤ 0.01 vs control). For panel D, E, F, H and G: Data are represented as mean ± SEM.

### Accumulation of p62 in ECs of patients with premature coronary artery disease and in the heart and vessels of Tchp knock-out mice

Accumulation of p62-positive aggregates are among the best-known characteristics of autophagy-deficient tissues (Komatsu et al, 2007). We analysed the expression of TCHP, p62 and cytokines in the ECs from patients with premature coronary artery disease (CAD). The ECs were obtained from the vessel wall of patients with endothelial dysfunction, comprising significant impairments in proliferation, adhesion and migration (Brittan et al, 2015). Gene expression analysis showed that ECs from patients express low level of TCHP and high level of p62 and cytokines in comparison with the ECs from healthy donors (Figure 7A). Treatment of ECs from patients with BAY11-7082 has reduced the accumulation of p62 puncta (Figure 7B) and cytokines expression (Figure 7C), whilst it improved their migratory capacity (Figure 7D). Finally, we analysed the accumulation of p62 in the tissue from Tchp knock-out mice and relative wild-type controls. p62 accumulated in the liver and pancreas of knock-out mice (**Figure S3**). Heart of Tchp knock-out mice presented a reduced cardiac vascularization with a significant accumulation of p62 in the cardiomyocytes and in the vessels (Figure 7E). Together, our data ascertains the strong link between depletion of TCHP, p62 and vascular function.

**Figure 7:**
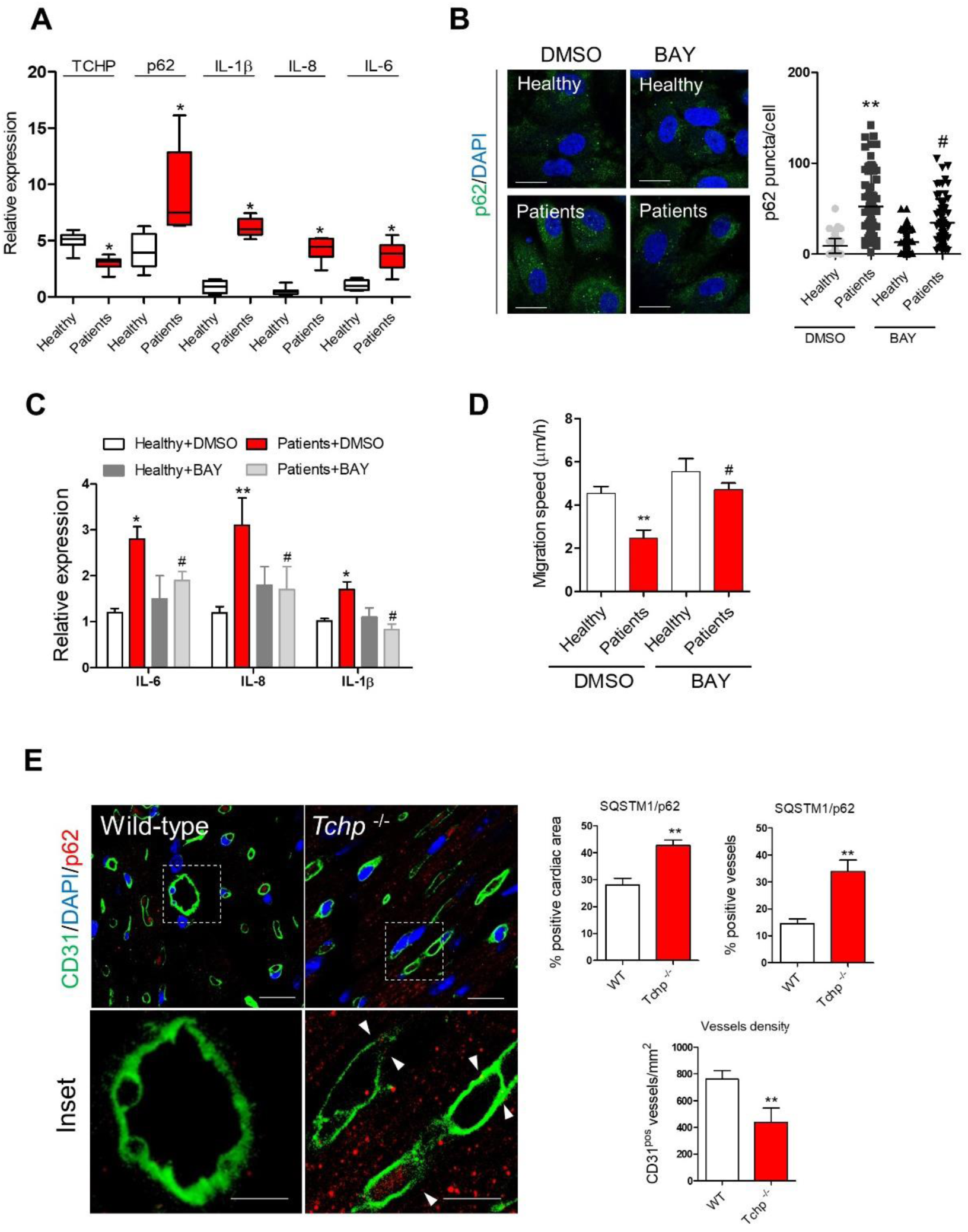
p62 accumulation in ECs from patients with CAD and Tchp knock-out mice. **A**, Expression of TCHP, p62, IL-1β, IL-8 and IL-6 in ECs from vessels wall from patients with CAD (n=8 per group, unpaired t test; *p<0.05 vs. Healthy subject). **B**, Staining and quantification of p62 in ECs from healthy subject and patients either treated with vehicle (DMSO) or BAY. Scale bar 50 µm; (n=3 per group; n=80 cells, n=3; unpaired t test; **p ≤ 0.01 vs control DMSO; ^#^p ≤ 0.05 vs shTCHP DMSO). **C**, Expression of IL-6, IL-8 and IL-1β and **D**, migration speed, in ECs from healthy subject and patients either treated with vehicle (DMSO) or BAY (300nM). (n=3; unpaired t test; *p ≤ 0.05; **p ≤ 0.01 vs control DMSO; ^#^p ≤ 0.05 vs shTCHP DMSO). **E**, *Left panels*: representative images of the heart of wild-type and Tchp knock-out mice stained for CD31 (green) and p62 (red). Scale bars, 100µm and 50µm for the inset *Right panels*: Quantification of percentage of cardiac area or vessels positive for p62 and vessel density. (n=4 per group; unpaired t test; **p ≤ 0.01 vs wild-type. Data are represented as mean ± SEM.

## Discussion

Accumulating evidence link the intact autophagic responses with the preservation of cardiovascular homeostasis in several physiological and pathological settings (Galluzzi et al, 2017b). The current work provides mechanistic insight into the action of TCHP on EC function, through impaired autophagy and accumulation of p62. Moreover, the block of autophagic flux in ECs resulted in the transcription of pro-inflammatory cytokines and accumulation of unfolded protein aggregates. Such phenotype is acquired through the activation of NF-kB pathway and could be rescued using an IKK inhibitor. A similar phenotype has been observed in endothelia specific ATG5 knock-out mice (Vion et al., 2017). These mice fed with high-fat diet developed endothelial dysfunction, vascular inflammation and atherosclerotic plaque (Vion et al, 2017).

Endogenous or expressed TCHP showed a dynamic localization in cells that could be due to a different repositioning of TCHP during different cell cycle stages or under the effect of cellular stressor and/or stimuli (Inaba et al, 2016; Nishizawa et al, 2005). Moreover, THCP localization in different subcellular compartments may mirror different functional roles played by the same protein. By immunoprecipitation and immunocytochemistry analysis, we demonstrated for the first time that TCHP binds PCM1 and co-localizes with PCM1 in pericentriolar satellites. Interestingly, TCHP depletion accelerated PCM1 proteasomal degradation. Although PCM1 depletion did not affect autophagy per se (Tang et al, 2013), it has destabilized GABARAP, but not LC3, through proteasomal degradation (Joachim et al, 2017).

GABARAP protein is important for maturation (Weidberg et al, 2010) and autophagosome fusion with lysosomes (Nguyen et al, 2016). We revealed that TCHP controls the stability of PCM1 and consequently of GABARAP, and this impacts on the maturation of the autophagosome. Using several approaches, we demonstrated that TCHP downregulation impairs basal autophagy leading to the accumulation of unresolved autophagosomes. Despite the TCHP depleted cells showing widespread alteration of autophagic flux; we still observed a strong activation of the endo-lysosomal pathway. Nevertheless, the autophagic flux is not completely blocked, but reversed by starvation, suggesting that TCHP-dependent reduction of basal autophagy is reversible and could be pharmacologically re-activated.

Another major finding in the present study is that TCHP depletion in cells and knock-out mice showed accumulation of p62, due to simultaneous reduction in autophagic flux and p62 transcriptional activation. An NF-kB response element has been identified in the p62 promoter (Vadlamudi & Shin, 1998). Our work showed, in agreement with previous studies (Huang et al, 2016), that NF-kB/p65 binds p62 promoter and plays an important role for p62 transcriptional regulation.

P62 was originally described as a scaffold protein and a signalling hub for the interactions with many types of enzymes through different binding domains (Katsuragi et al, 2015). It is well-known also that p62 induces inflammatory cytokines production via TRAF6 polyubiquitination and thereby NF-κB activation (Duran et al, 2008). Instead, a previous study demonstrated that stimulated autophagy, by enhanced degradation of p62, reduced inflammation, whereas blocking autophagy had an opposite effect (Shi et al, 2012). Thus far, the phenotypic screens identified genes which reduce the accumulation of p62 following stress stimuli, as a novel approach to map autophagy pathways (DeJesus et al, 2016; Strohecker et al, 2015). While our screen was designed to identify compounds targeting the accumulation of p62, it is likely that amongst our hits we also identified compound which regulate autophagy. Being that NF-kB is a critical mediator of endothelial cell dysfunction and impairs vascular regeneration (Caporali et al, 2015), reactivation of autophagy through inhibition of NF-kB may help to restore vascular function and reparative angiogenesis. In conclusion, these results reveal for the first time the pivotal role for TCHP in linking EC function with the control of autophagy, highlighting a novel possible role in vascular disease.

## Material and Methods

### Cells and cell culture and reagents

Human Umbilical Vein ECs (HUVECs) and ECs from healthy donor and patients were cultured in EGM-2 (EBM-2 + SingleQuots™ Kit) and 2% Foetal Bovine Serum (FBS) (Lonza). HUVECs and ECs were used between P2 and P6 passage. HEK293T were grown in Dulbecco’s modified eagle’s medium (DMEM) with 4.5g/L glucose, 2mM L-Glutamine, without Na Pyruvate (Lonza), 10% FBS and 1% Penicillin/Streptomycin (Pen/Strep). *Reagents*: Bafilomycin A1 (CST), Torin-1 (CST), BAY11-7082 (Abcam), TYRPHOSTIN AG1288 (Abcam), SB202190 (Tocris).

### Lentiviral vectors and plasmid constructs

The pLKO DNA plasmids containing the shRNA sequence against human TCHP, was purchased from Sigma Aldrich (Mission®RNAi TRCN0000127662 and TRCN TRCN0000130868). The scrambled sequence shRNA plasmid was purchased from Addgene, plasmid #1864. The packing plasmids used were pCMV-dR8.2 dvpr, plasmid #8455 and pCMV-VSVG, plasmid #8454 from Addgene. p62-V5 (HsCD00434166), TCHP-V5 (HsCD00444989) and empty vector (pLX304) are from DNASU Plasmid collection, pBABE-puro mCherry-EGFP-LC3B was a gift from Jayanta Debnath (Addgene plasmid # 22418). To generate an expression plasmid for 3xFLAG–tagged, the full-length TCHP coding sequence was amplified by PCR with primers having the sequence 5’-GATGACAAGCTTGGAAACTCCGAGCCTCAGAGA-3’ and 5’ GGATCCTCTAGATTCTCTGTACTTATGGTACCC-3’. The PCR product was digested with HindIII and XbaI and the resulting DNA fragment was inserted into p3xFLAG-CMV-7.1 (Sigma) to prepare p3xFLAG-TCHP. A 3xFLAG–TCHP coding sequence was then amplified by PCR, the products were gel purified and verified by sequencing. The deletion mutants FLAG-TCHP Δ1, lacking the first coiled motif of TCHP protein, and FLAG-TCHP Δ1.2, lacking the first and second coiled-coil motives, were generated by PCR starting from the previously described p3XFLAG-TCHP full length construct. The products of the PCR were gel purified, verified by sequencing, and cloned into the HindIII-XbaI sites of p3x-FLAG-CMV-7.1 expression vector.

The primers used in PCR are: TCHP Δ1 Fw_: 5’-GCGATTAAGCTTTTCAGGATGTCTGACATCTGC–3’; TCHP Δ1.2 Fw_: 5’-GCGATTAAGCTTCAACTTTTGTACGAACACTGG–3’ and TCHP Rw_: 5’-GGATCCTCTAGATTCTCTGTACTTATGGTACCC–3’

### Endothelial cells functional assays

The following functional assays were performed: Matrigel assay with HUVECs was performed as previously described using BD Matrigel Basement Membrane Matrix (BD Biosciences). Migration was analyzed with the ECIS chip array (8W1E) (Applied Biophysics). The migration speed was calculated in micrometers per hour.

### Proteinase K protection assay

The subcellular fractionation (PNS; postnuclear supernatant = 300 × g for 5 min at 4°C, LSP; low-speed pellet = 700 × g for 5 min at 4°C, HSP; high-speed pellet, HSS; high-speed supernatant = 100,000 × g for 30 min at 4°C) of control of TCHP knock-down ECs was performed as described in (Hasegawa et al, 2016). In brief, each fraction of LSP and HSP was treated with 100 μg/ml proteinase K on ice with or without 0.5% Triton X-100 for 30 min. The fraction samples were precipitated with 10% trichloroacetic acid, washed with ice-cold acetone three times, resuspended in sample buffer including 3 M urea, and then analysed by Western blot for p62, NDP52 and GAPDH antibody. GAPDH was used as loading control as described in (Hasegawa et al, 2016).

### In Vivo Matrigel Plug Assay

Experiments involving mice were covered by project and personal licenses issued by the UK Home Office, and they were performed in accordance with the Guide for the Care and Use of Laboratory Animals (the Institute of Laboratory Animal Resources, 1996) and in accordance with Animal Research Report of In vivo Experiments (ARRIVE) guidelines. CD-1 mice (male, 10 weeks old) were subcutaneously injected into the groin regions with 400 μL Matrigel containing recombinant mouse basic FGF (PeproTech, 250 ng/mL) and heparin (Sigma, 50 U/mL) mixed with control or TCHP siRNA (Dharmacon) (lipids [Lipofectamine RNAiMAX reagent, ratio 1:1 in volume] 5 μg/gel, n = 5 per group). After 21 days, mice were sacrificed, and the Matrigel plugs were removed and fixed in 4% paraformaldehyde.

### Immunohistochemistry

Paraffin cross-sections were blocked with normal goat serum, incubated with anti-CD31(Abcam; 1:200) and anti-p62 (Genetex 1:200) primary antibody overnight at 4°C, and then incubated with Alexa 488-conjugated anti-rat IgG antibody (Thermo Fisher). Microvessel density was quantified in 10 fields/section. CD31-positive vessel area was quantified using ImageJ software and expressed per square micrometres.

### RNA extraction and quantitative real-time analysis

Total RNA was extracted using miReasy kit (Qiagen). For mRNA analysis, cDNA was amplified by quantitative real-time PCR (qPCR) and normalized to 18S ribosomal RNA. Each reaction was performed in triplicate. Quantification was performed by the 2^−ΔΔCt^ method (Schmittgen & Livak, 2008). Primers are from Sigma (KiCqStart^TM^ Primers).

### Chromatin Immunoprecipitation

Nuclei were isolated from formaldehyde (1% final)-fixed HUVECs by lysing in ChIP Lysis Buffer (1% SDS, 10 mM EDTA, 50 mM Tris-HCL [pH 8.1]) supplemented with protease inhibitors. Chromatin was fragmented by sonication using a Bioruptor UCD-300 ultrasound sonicator (Diagenode). DNA-cross-linked proteins were immunoprecipitated (1% kept as input) using 5 μg of NF-kB/p65 (Millipore) or control mouse IgG antibody. The antibody was pulled down with protein G beads (Dynabeads, Thermo Fishers) at 4°C overnight. Associated DNA was then purified by extraction using Monarch PCR & DNA Cleanup Kit (New England Biolabs). Immunoprecipitated DNA and total input were used as a template for real-time qPCR. The ChIP primers for NF-kB/p65 on IkBα promoter are: Fw GTGCGCCCTCAACTAACAGT Rev CATCCCAATGAAGCTTCTGA. Identification of the NF-kB/p65 binding sites in the p62 promoter was performed retrieving ENCODE dataset GSM935527 (Chr 5: 179,245,409-179,256,832). Primers for NF-kB/p65 on p62 promoter are: Fw CTAAAGATGGCCCAGAGCAG Rev CCCCCTCCCAAATAATCCTA.

### Mass Spectrometric analysis

Gel bands were subjected to overnight trypsin digestion and peptide extracts were dried by Speedvac. The dried peptide samples were re-suspended in MS-loading buffer (0.05% trifluoroacetic acid in water) and then filtered using Millex filter before HPLC-MS analysis. Nano-ESI-HPLC-MS/MS analysis was performed using an online system consisting of a nano-pump (Dionex Ultimate 3000, Thermo Fisher) coupled to a QExactive instrument (Thermo Fisher). MS/MS Fragmentation was performed under Nitrogen gas using high energy collision dissociation in the HCD cell. Data was acquired using Xcalibur ver 3.1.66.10. Data from MS/MS spectra was searched using MASCOT Versions 2.4 (Matrix Science Ltd) against the Human subset of Uniprot database with maximum missed-cut value set to 2. Differentially expressed proteins were considered significant if the p-value was less than 0.05 and if the number of peptides used in quantitation per protein was equal to or more than 2.

### Phenotypic screening assay

#### Image Acquisition

Plates were imaged on a wide-field Imagexpress Micro XL high content microscope (Molecular Devices). Images of Hoechst labelled nuclei and p62 antibody labelling were imaged using the DAPI and Cy3 filter sets from 4 different sites within the well using a 20X S Plan Fluor objective containing up to 200 cells per field of view. *Image Analysis:* Images were analysed using a custom workflow developed in the MetaXpress Custom Module Editor (Molecular Devices). *Data analysis:* Data handling and analysis was done using Spotfire High Content Analyser software (PerkinElmer).

### Western blot analyses

Total proteins were extracted in RIPA buffer containing 1 mM sodium orthovanadate and Complete Protease Inhibitor Cocktail (Roche Applied Science) and quantified using the Pierce™ BCA protein assay kit (Thermo Fisher). Equal amounts of proteins were loaded onto SDS-Polyacrilamide gels and transferred to PVDF membrane. The membranes were then blocked with 5% non-fat milk in TBST 0.1% and immunoblotted overnight at 4°C with the following primary antibodies: TCHP (Santa Cruz Biotechnology, SC-515025), β-actin (Sigma, A5441), p62 (GeneTex, GTX100685), LC3 (CST, #2775), p16 (BD Bioscience, 551154), NF-kB (Milllipore, MAB3026), (S536) NF-kB (CST, #3033), PCM1 (CST, #5213) and GABARAP (CST, #26632) and NDP52 (. Secondary antibodies: anti-Mouse IgG–Peroxidase (Sigma, A5906), anti-Rabbit IgG–Peroxidase (Sigma, A0545) were incubated for 1 hr at RT. Pixel intensity/quantification was performed using ImageJ.

### Immunofluorescence

HUVECs cells were plated on fibronectin-coated glass coverslips. Twenty-four hours later, the slides were fixed with 4% paraformaldehyde, permeabilized with 0.05% Triton X-100 in PBS, then incubated with primary antibody in 3% BSA overnight at 4^ο^ C. Secondary antibodies diluted 1:1000 in 3% BSA. Slides were imaged on Zeiss LSM-780 confocal. Primary and secondary antibodies used for immunofluorescence were: PCM1 (CST, #5213), p62 (GeneTex, GTX100685), LC3 (CST, 2775), Rab7 (CST, 9367), Rab11 (CST, 5589), EEA1 (CST, 3288) and LAMP2 (CST, 49067). Aggregates of ECs was measured using the PROTEOSTAT kit (Enzo Life Science, ENZ-51035-0025). Senescence of ECs was measured using the Cellular Senescence Assay kit from Cell Biolabs, Inc. (Cat: CBA-230). The image analysis software Cell Profiler was used to quantify co-localization.

### Electron microscopic analysis

Cell pellets were prepared and fixed in 4% paraformaldehyde or 2.5% paraformaldehyde/0.2% glutaraldehyde in 0.1 M phosphate buffer (pH 7). Pellets were embedded in HM20, 70 nm sections cut and examined using a Philips CM10 transmission electron microscope equipped with a Gatan Bioscan Camera.

### Endothelial cells from patients

The study was performed with the approval of the South-East Scotland Research Ethics Committee, in accordance with the Declaration of Helsinki and with the written informed consent of all participants. Patients with premature coronary artery disease and a family history of premature coronary artery disease (n = 8) were identified from the outpatient department, Royal Infirmary of Edinburgh, Scotland, UK. A control group of healthy age- and sex-matched subjects (n = 8) with no evidence of significant coronary artery disease following computed tomography coronary angiography (CTCA) was recruited from the Clinical Research Imaging Centre, Royal Infirmary of Edinburgh. Subjects attended the Clinical Research Facility at the Royal Infirmary of Edinburgh for vascular assessment and tissue sampling. Vessel wall endothelial cells were isolated by wire biopsy for in vitro expansion as reported in (Brittan et al, 2015).

### TCHP knock-out mice tissues

Heart tissues are from C57BL/6N-Atm1Brd Tchp^tm1b (EUCOMM)Wtsi/WtsiIeg^ (Tchp knock-out) mice and were kindly provided from the Wellcome Trust Sanger Institute. Histological analysis was performed on 4 wild-type mice (2 male and 2 female) and 4 Tchp knock-out mice (2 male and 2 female) 16-week-old.

### Statistical analysis

Comparisons between different conditions were assessed using 2-tailed Student’s *t* test. If the normality test failed, the Mann-Whitney test was performed. Continuous data are expressed as mean ± SEM of three independent experiments. *P* value <0.05 was considered statistically significant. Analyses were performed using GraphPad Prism v5.01.

## Acknowledgments

We acknowledge the assistance of the microscopy and histology core facility at the University of Aberdeen and Lorraine Rose for technical help at University of Edinburgh. This study was supported by grants from British Heart Foundation (BHF) (FS/11/52/29018) for A.C. and (FS/14/7/30574) for A.M; A.C. and N.O.C. acknowledge the support of the Wellcome Trust-University of Edinburgh Institutional Strategic Support Fund and MRC-IMPC Pump Priming Award (MR/R014353/1). T.M is supported by BHF Career Re-entry Fellow (FS/16/38/32351); M.B. is a BHF Intermediate Fellow (FS/16/4/31831). A.C. acknowledges Edinburgh BHF Research Excellence Award.

## Authors Contributions

A.M. acquired and analysed the data and drafted the manuscript; A.L.; D.DA; M.V.; N.G., T.M.; D.M.; E.P.; M.B. N.M. and J.C.D participated in acquiring and analyses of the data and revised the manuscript; N.O.C and J.C.D. designed the phenotypic screening assays and high content data analysis protocols and contributed to revising the manuscript; L.E. performed the mass spectrometry experiments; A.C. designed the overall study and drafted the manuscript.

