## Supplemental Figures and Material for "Trichoplein controls endothelial cell function by regulating autophagy"

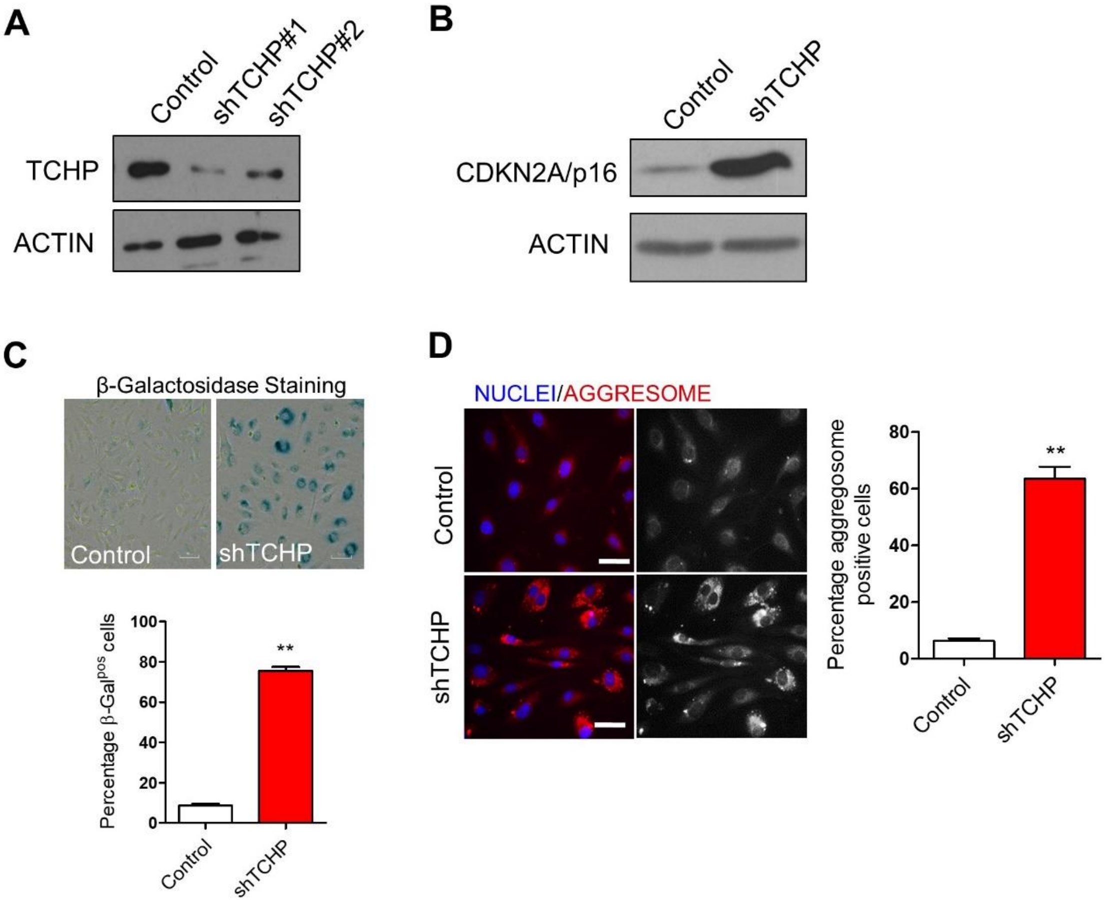


**Figure S1: Premature senescent phenotype in TCHP knock-down cells**

**A,** Western blot anti-TCHP following knock-down of TCHP, B**,** Western Blot anti-p16 **C**, β-Galactosidase activity as reveal by the chromogenic β-Gal substrate X-Gal. Scale bar 25µm. (n= 5, unpaired t test, **p ≤ 0.01 vs control). **E,** Aggresome staining and quantification of protein aggregates; Scale bar 25µm. Data are mean ± SEM, (n= 5, unpaired t test, **p ≤ 0.01 vs control)

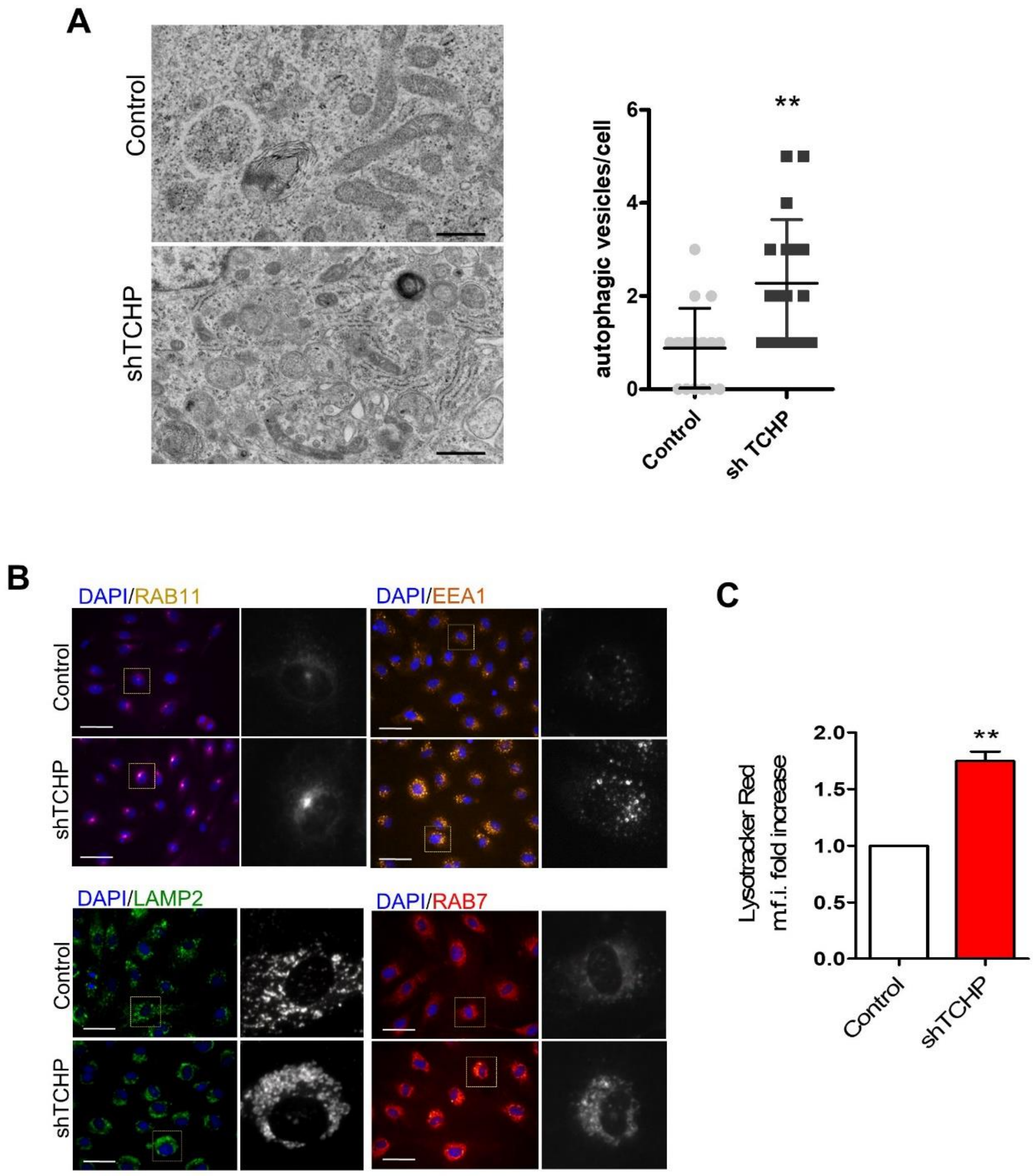


**Figure S2: Analysis of autophagic features in TCHP-depleted endothelial cells**
**A,** Representative pictures from Transmission Electron Microscopy analysis. Scale bar 500nm. Left panel: quantification of autophagic vacuoles . Data are mean ± SEM (n=16, unpaired t test; **p ≤ 0.01 vs control). **C**, Immunofluorescence for recycling (RAB11), early (EEA1), late (RAB7) endosomes and lysosome (LAMP1/2); Scale bar 25µm; **D**, Quantitative analysis of Lysotracker Red by flow cytometry. Data are mean ± SEM (n=3, unpaired t test; **p ≤ 0.01 vs control).

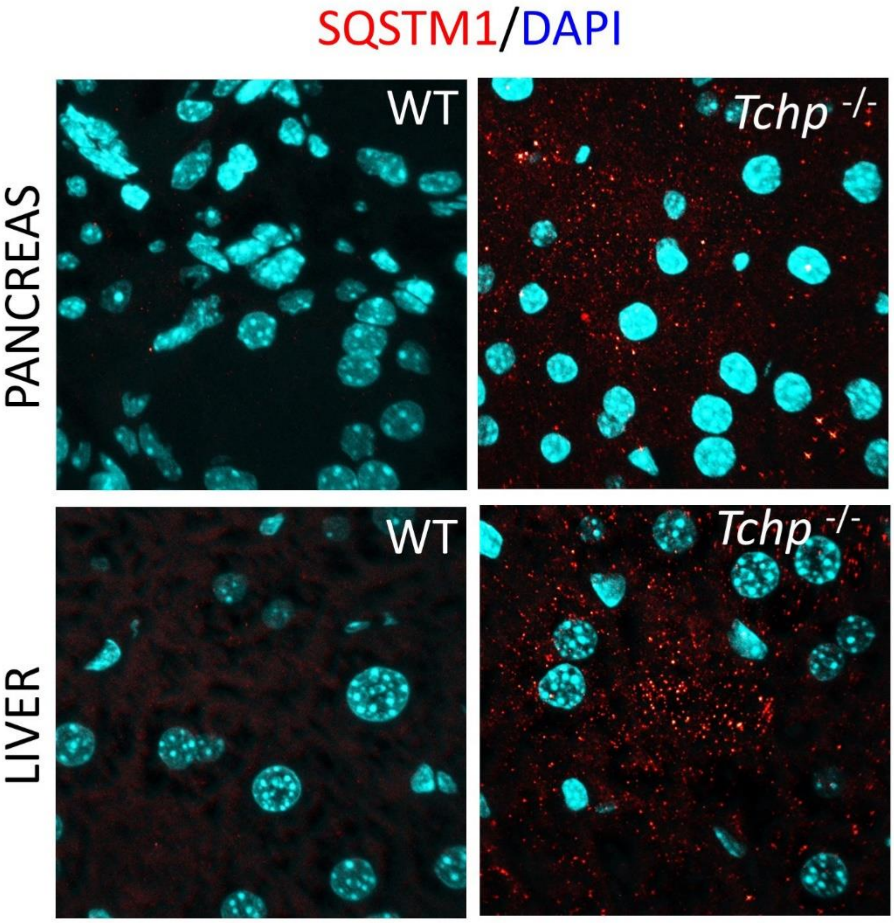


**Figure S3:** Representative images of the heart of wild-type and Tchp knock-out mice stained for DAPI (blue) and p62 (red). Scale bars, 100µm

### Expanded Methods

#### Phenotypic screening assay

**Image Acquisition:** Plates were imaged on a wide-field Imagexpress Micro XL high content microscope (Molecular Devices). Images of hoechst labelled nuclei and p62 antibody labelling were imaged using the DAPI and Cy3 filter sets from 4 different sites within the well using a 20X S Plan Fluor objective containing up to 200 cells per field of view. **Image Analysis:** Images were analysed using a custom workflow developed in the MetaXpress Custom Module Editor (Molecular Devices). A top hat filter (size = 10 pixels, shape = circle) was applied to remove background fluorescence from the p62 images. Using the granularity module, nuclei were first identified in the DAPI image using a user-defined intensity above local threshold method and maximum (30 µm) and minimum (10 µm) widths. p62 puncta was then detected by using a user-defined intensity above local background and maximum (1 µm) and minimum (0.5 µm) widths. The nuclear objects were then used as seeds to create a pseudo-whole cell by growing the mask by 50 pixels. Finally, the nuclear region was subtracted from the whole cell mask to give nuclear and cytoplasmic masks. The number of p62 puncta in the nuclear and cytoplasmic regions was counted per cell. Features measured are listed in provided Supplemental Table. **Data analysis:** Data handling and analysis was done using Spotfire High Content Analyser software (PerkinElmer). Data was aggregated to whole well averages and was plate normalised to the negative controls by dividing all values on the plate by the median value for the negative control and then scaling the values between 0 and infinity, with 1 being the median of the negative controls on that plate. For hit identification, control cells (no shRNA) were used as positive controls and an ensemble-based tree classifier was used to identify hit compounds with a cross validated 5-fold CV mis-classification error of 0.4%. The following features were used in the ensemble-based tree classifier: Cell_Count; Nuclear_Granules_Nuclear_count_Sum; Cytoplasmic_Granules_Nuclear_Count_Sum.

**Cell Profiler Workflow for co-localization**

The image analysis software Cell Profiler was used to quantify co-localisation within the images obtained by confocal microscopy. The images were processed and subjected to a 20-module co-localisation workflow. To summarise, the images are initially loaded as sperate channels, corrected for light illumination and aligned. The images then undergo pixel-based correlation, where, the pixel intensities are compared in each image and any initial correlation determined. The specific structures of interest are determined by thresholding the images and segmenting them into objects, this allows for any co-localisation to be determined between the individual channels and objects of interest. During the final stage of image analysis, the images are further enhanced, and the objects of interest are refined. The software then calculates various statistics within the defined regions and calculates the number of pixels within a specific object in each channel. The area occupied by co-localised regions is then divided by the area of co-localised objects of interest and a per image pixel fraction determined from the total object pixels. This information is then converted into a percentage co-localisation and the data exported. A detailed description of each stage of the Cell Profiler workflow is located on their website (http://cellprofiler.org).

**Mass Spectrometric analysis**

Gel bands were subjected to overnight trypsin digestion and peptide extracts were dried by Speedvac. The dried peptide samples were re-suspended in MS-loading buffer (0.05% trifluoroacetic acid in water) and then filtered using Millex filter before HPLC-MS analysis. Nano-ESI-HPLC-MS/MS analysis was performed using an on-line system consisting of a nano-pump (Dionex Ultimate 3000, Thermo-Fisher, UK) coupled to a QExactive instrument (Thermo-Fisher, UK) with a pre-column of 300 µm x 5 mm (Acclaim Pepmap, 5 µm particle size) connected to a column of 75 µm x 50 cm (Acclaim Pepmap, 3 µm particle size). Samples were analysed on a 90 min gradient in data dependent analysis (1 survey scan at 70k resolution followed by the top 10 MS/MS). The gradient between solvent A (2%Acetonitrile in water 0.1%formic acid) and solvent B (80% acetonitrile-20% water and 0.1% formic acid) was as follows: 7min with buffer A, over 1 min increase to 4% buffer B, 57min increase to 25% buffer B, over 4min increase to 35%, over 1 min increase to 98% buffer B and stay under those conditions for 9min, switch to 2% buffer B over 1 min and the column was conditioned for 10min under those final conditions. MSMS Fragmentation was performed under Nitrogen gas using high energy collision dissociation in the HCD cell. Data was acquired using Xcalibur ver 3.1.66.10. Data from MS/MS spectra was searched using MASCOT Versions 2.4 (Matrix Science Ltd, UK) against the Human subset of Uniprot database with maximum missed-cut value set to 2. Following features were used in all searches: i) variable methionine oxidation, ii) fixed cysteine carbamidomethylation, iii) precursor mass tolerance of 10 ppm, iv) MS/MS tolerance of 0.05 amu, v) significance threshold (p) below 0.05 (MudPIT scoring) and vi) final peptide score of 20. Progenesis (version 4 Nonlinear Dynamics, UK) was used for LC-MS label-free quantitation. Only MS/MS peaks with a charge of 2+, 3+ or 4+ were taken into account for the total number of ‘Feature’ (signal at one particular retention time and m/z) and only the five most intense spectra per ‘Feature’ were included. Results were exported using a peptide score cut off of 20. From the exported results sheet, differentially expressed proteins were considered significant if the p-value was less than 0.05 and if the number of peptides used in quantitation per protein was equal to or more than 2.
